# Methodological guidelines for isolation and purification of plant extracellular vesicles

**DOI:** 10.1101/2021.09.01.458648

**Authors:** Yifan Huang, Shumei Wang, Qiang Cai, Hailing Jin

**Affiliations:** State Key Laboratory of Hybrid Rice, College of Life Science, Wuhan University, Wuhan, 430072, China; Department of Microbiology and Plant Pathology and Center for Plant Cell Biology, Institute for Integrative Genome Biology, University of California, Riverside, California 92507, USA; Hubei Hongshan Laboratory, Wuhan, 430072, China

## Abstract

Plant extracellular vesicles (EVs) have become the focus of rising interest due to their important roles in the cross-kingdom trafficking of molecules from hosts to interacting microbes to modulate pathogen virulence. However, the isolation of pure intact EVs from plants still represents a considerable challenge. Currently, plant EVs have been isolated from apoplastic washing fluid (AWF) using a variety of methods. Here, we compare two published methods used for isolating plant EVs, and provide a detailed recommended method for AWF collection from *Arabidopsis thaliana*, followed by EV isolation via differential ultracentrifugation. To further separate and purify specific subclasses of EV from heterogeneous vesicles, sucrose or iodixanol density-based separation and immunoaffinity capture are then utilized. We found that immunoaffinity capture provides a significant advantage for specific EV isolation when suitable specific EV biomarkers and their corresponding antibodies are available. Overall, this study guides the selection and optimization of EV isolation methods for desired downstream applications.

## INTRODUCTION

Cell-to-cell communication between plants and pathogens requires the secretion and delivery of molecular signals into extracellular environments and their subsequent transport into interacting organisms. This process is critical for the survival of plants and pathogens (Kimura et al. 2001; Mahlapuu et al. 2016; Toruno et al. 2016). Recent studies have demonstrated that RNAs, including regulatory small RNAs (sRNAs), are able to move between pathogens and their hosts and regulate biological processes in recipient cells (Knip et al. 2014; Zhang et al. 2016b; Cai et al. 2018; Cai et al. 2019b; Huang et al. 2019). For a long time, the mechanisms used to pass these sRNAs through multiple barriers and into the opposing host or fungal cells were largely unknown. However, recent studies have shown that extracellular vesicles (EVs) can traffic sRNAs from plants to their pathogens (Cai et al. 2018). Currently, plant EVs have generated further interest because of their numerous functions in bioactive molecule exchange and cell-to-cell communication (Mathieu et al. 2019; Cai et al. 2021; Kameli et al. 2021).

EVs are small, lipid bilayer-enclosed vesicles containing various protein and RNA cargoes, and are released by cells of eukaryotic and prokaryotic organisms into the extracellular space (Colombo et al. 2014; van Niel et al. 2018). EVs are a heterogeneous group of vesicles with different sizes and intracellular origins. They are classified into exosomes, microvesicles and apoptotic bodies, which originate from the multivesicular bodies (MVBs), shed from the plasma membrane, or an apoptotic cell during apoptosis, respectively (Akers et al. 2013; Colombo et al. 2014; van Niel et al. 2018). In plants, EVs were initially observed in carrot cell cultures by transmission electron microscopy (TEM) in 1967 (Halperin and Jensen 1967). Since then, plant EVs have been observed to be enriched in fungus-plants interaction sites by TEM, such as *Blumeria graminis f. sp*.*hordei* infected barley epidermal cells (An et al. 2006a; An et al. 2006b), *Botrytis cinerea* infected *Arabidopsis* leaf cells (Cai et al. 2018), and *Rhizophagus irregularis* arbuscules in rice root (Roth et al. 2019). TEM and confocal microscopy analysis demonstrated that plant MVBs fused with the plasma membrane underlie fungal or oomycete invasion sites, suggesting that plant exosomes are released by MVB mediated secretion (An et al. 2006a; An et al. 2006b; An et al. 2007; Nielsen et al. 2012; Bozkurt et al. 2014; Cai et al. 2018).

It is worth noting that plant EVs are nanovesicles primarily from the apoplastic washing fluid (AWF). However, nanovesicles isolated from disrupted whole leaf tissue are not pure EVs, as they contain cytoplasmic intracellular membrane contaminants (Liu et al. 2020). Currently, plant EVs have been isolated from the AWF of a few different plant tissues, including *Arabidopsis* leaves (Rutter and Innes 2017; Cai et al. 2018; He et al. 2021), sunflower seeds and seedlings (Regente et al. 2009; Regente et al. 2017), olive pollen tubes (Prado et al. 2014) and *Nicotiana benthamiana* leaves (Movahed et al. 2019). In *Arabidopsis* leaves, to our knowledge, at least three known EV subtypes exist: Tetraspanin (TET) 8-positive EVs derived from MVBs which can be considered bona fide plant exosomes (Cai et al. 2018; Cai et al. 2021), Penetration 1 (PEN1)-positive EVs (Rutter and Innes 2017), and EVs produced by, exocyst-positive organelle (EXPO)’s fusion with the plasma membrane (Wang et al. 2010; Ding et al. 2014). It has been demonstrated that plant endogenous sRNAs are secreted by EVs as a defense mechanism against fungal pathogens (Cai et al. 2018). Further, an additional study demonstrated that TET8-positive exosomes are the major class of plant EVs that transport sRNAs, along with several RNA binding proteins which contribute to sRNA selective loading and stabilization in EVs (He et al. 2021).

In animals, many isolation methods for EVs have been developed in the last decade. Of these methods, differential ultracentrifugation separation is classically considered the standard for EV isolation, specifically for the isolation of small EVs or exosomes (Thery et al. 2006; Mathivanan et al. 2012). This method has several substeps, including centrifugation at 300 ×g to sediment cells, at 2000 ×g to remove dead cells and apoptotic bodies (large vesicles), at 10,000-15,000 ×g to remove cell debris and microvesicles (medium vesicles), and a final centrifugation step at ≥ 100,000 ×g (100,000 to 200,000 ×g) to pellet small EVs and exosomes (Thery et al. 2006; Crescitelli et al. 2013; Konoshenko et al. 2018; Willms et al. 2018; Jeppesen et al. 2019). Then, the EV pellet is washed once to remove non-EV proteins by resuspension and the final centrifugation step is repeated (Thery et al. 2006; Konoshenko et al. 2018). The pellet produced by differential ultracentrifugation can be further separated by extra steps, such as high-speed density gradient ultracentrifugation or bead-based immunoaffinity capture, which leads to the isolation of subtypes of EVs and increases the purity of isolated EVs (Thery et al. 2006; Jeppesen et al. 2019).

While animal EVs have been well studied over the last decades, plant EVs remain poorly investigated. This is mainly due to the lack of accepted EV isolation protocols. Because plant EVs are derived primarily from the apoplastic space, the crucial first step to EV isolation is isolation of the AWF, which is obtained by a simple, well-established infiltration-centrifugation method (Wang et al. 2005; Sanmartin et al. 2007; Hatsugai et al. 2009; O’Leary et al. 2014). Based on established animal EV purification protocols, plant EV separation involves differential ultracentrifugation of AWF, with two consecutive steps of low speed centrifugation at 2,000 ×g and 10,000 ×g to remove dead cells, cell debris and large vesicles, and high speed centrifugation at 100,000 × g to pellet the small EVs (Prado et al. 2014; Cai et al. 2018; Movahed et al. 2019; Liu et al. 2020; He et al. 2021). In some studies, the lower centrifugal force, 40,000 ×g was used to generate a final pellet from the EV fractions, which contained PEN1-positive EVs in *Arabidopsis* (Rutter and Innes 2017) and EVs derived from sunflower seeds and seedlings (Regente et al. 2009; Regente et al. 2017). Note that in distinct protocols for plant EV isolation, the differences lie not only in the speed of centrifugation for the final EV sedimentation, but also in how the AWF is collected. So far, there is still no standard protocol for plant EV isolation from AWF across different plant species. Thus, in this study, we compared the results from different methods, propose a standardized method for plant EV isolation and purification from *Arabidopsis*. We performed high-speed density gradient ultracentrifugation that enables distinguishing EV subtypes floated in an iodixanol or sucrose gradient to be separated and purified based on their different densities. Furthermore, we also describe a recently developed immunoaffinity capture method, using bead-based antibodies that recognize the plant EV enriched TET8 protein, allowing the precise capture of the specific TET8-positive EV subtype.

## RESULTS

### Isolation of plant EVs by differential ultracentrifugation

Differential ultracentrifugation is the most commonly used method for EV isolation from cell culture supernatants and biological fluids. The detailed protocol was published by Théry *et al*. in 2006 (Thery et al. 2006; Willms et al. 2018). The methods used to isolate plant EVs share similarities with that used for mammalian EVs isolation, with the additional first step of collecting AWF from leaves (Figure 1). Based on the commonly used infiltration-centrifugation method for plant AWF collection, we developed a protocol for the extraction of AWF from *Arabidopsis* leaves with an optimized vacuum infiltration and centrifugation method (O’Leary et al. 2014; Cai et al. 2018; He et al. 2021). Initially, fully expanded rosette leaves were detached from plants at the base of the leaf using a razor blade to limit as much the phloem stream as possible, which contains mobile RNAs and ribonucleoprotein complexes (Zhang et al. 2009; Liu and Chen 2018). Cytoplasmic contaminants from damaged cells were removed by washing the cut leaves in distilled water (Figure 1A). The detached leaves were infiltrated with the infiltration buffer by negative pressure within a needleless syringe, then the leaves were carefully arranged to have the base of the leaves with the cutting site towards the bottom of the tube to avoid cell damage during centrifugation (Figure 1A). AWF was then collected by centrifugation at 900 ×g (Figure 1A). EVs were isolated from AWF by the following centrifugation steps (Figure 1B): (i) The AWF was centrifuged for 30 min at 4 °C at 2,000 × g to remove large cell debris. (ii) The supernatants were filtered through a 0.45 μm filter to remove the largest vesicles. (iii) The supernatant was moved into new ultracentrifuge tubes, and large vesicles were removed with another centrifugation step at 10,000 × g for 30 minutes at 4°C. (iv) The plant EV fraction was then pelleted at a high speed (100,000 × g) centrifugation for 1 hour. (v) The pellet was washed to remove potential protein aggregates via a second round of ultracentrifugation at 100,000 × g. The EV pellet of this step is called the P100 fraction.

**Figure 1.**
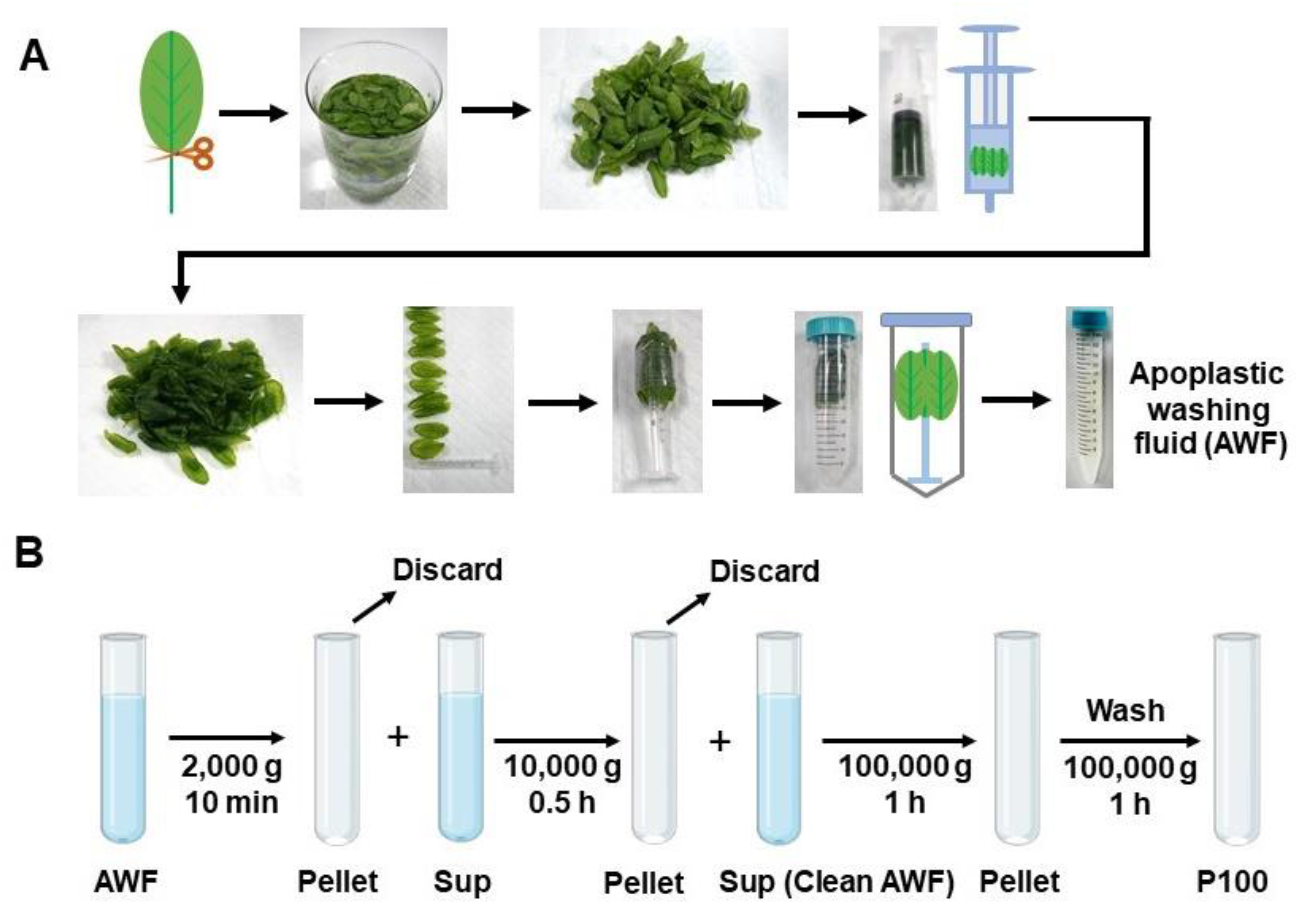
Schematic of the plant EVs isolation workflow. (A) Images show the various steps in apoplastic washing fluid (AWF) isolation from *Arabidopsis* (detached leaves protocol, Method 1 in Figure 2). The distinct proximal (petiole) part of leaves was removed using scissors, and the distal (blade) zones of leaves were collected. The leaves were placed in a syringe and gently vacuumed with infiltration buffer. The syringe with taped leaves was placed into a 50 ml conical tube, and then centrifuged at 900 × g to collect the apoplastic washing fluid. (B) Schematic of EVs isolated by differential ultracentrifugation of AWF from *Arabidopsis*. The clean AWF was centrifuged at 100,000 × g to obtain the P100 EV fraction. Sup, Supernatant.

### Technical evaluation of AWF collection from *Arabidopsis* leaves

Extraction of AWF is the crucial step to obtain high quality plant EVs with few contaminants. In parallel with the detached leaves method described above (Figure 1A, Method 1), Rutter *et al*. harvested AWF from the whole aerial part of plants that were harvested by cutting off the root, and then were vacuumed and centrifuged to collect the AWF, referred as Method 2 in this paper (Figure S1) (Rutter and Innes 2017; Baldrich et al. 2019). Here, we compared Method 1 to Method 2 for purity and quality of EVs. Previously we showed that fungal infection increases EV secretion (Cai et al. 2018; He et al. 2021). Here, we used *B. cinerea*-infected *Arabidopsis* to increase the yield of isolated EVs. Ideally, AWF should be free of contamination by cell debris and cytoplasmic molecules, such as chlorophyll, the major pigment (green) in chloroplasts (O’Leary et al. 2014). However, AWF extracted by Method 2 was green in color indicating obvious contamination of cytoplasmic molecules, whereas the AWF extracted by Method 1 is clear with no visible chlorophyll contamination (Figure 2A). Western blot analysis showed that the AWF extracted by Method 2 had a much higher abundance of Rubisco proteins than the AWF sample extracted by Method 1, and the Rubisco protein was also present at a high level in the P100 sample derived from AWF extracted by Method 2 (Figure 2B). To be more precise, we directly visualized the vesicles in the P100 fractions prepared from AWF extracted by Method 1 and Method 2, using TEM. Much cleaner vesicles were observed in the P100 fraction obtained via Method 1 (Figure 2C). The P100 fraction obtained from Method 2 contains large amounts of non-vesicle structures/materials, while much less non-vesicle material was observed in the P100 fraction obtained via Method 1 (Figure 2C). These results indicate that the detached leaves method (Method 1) is a better choice for collecting AWF to reduce contamination in EV preparations.

**Figure 2.**
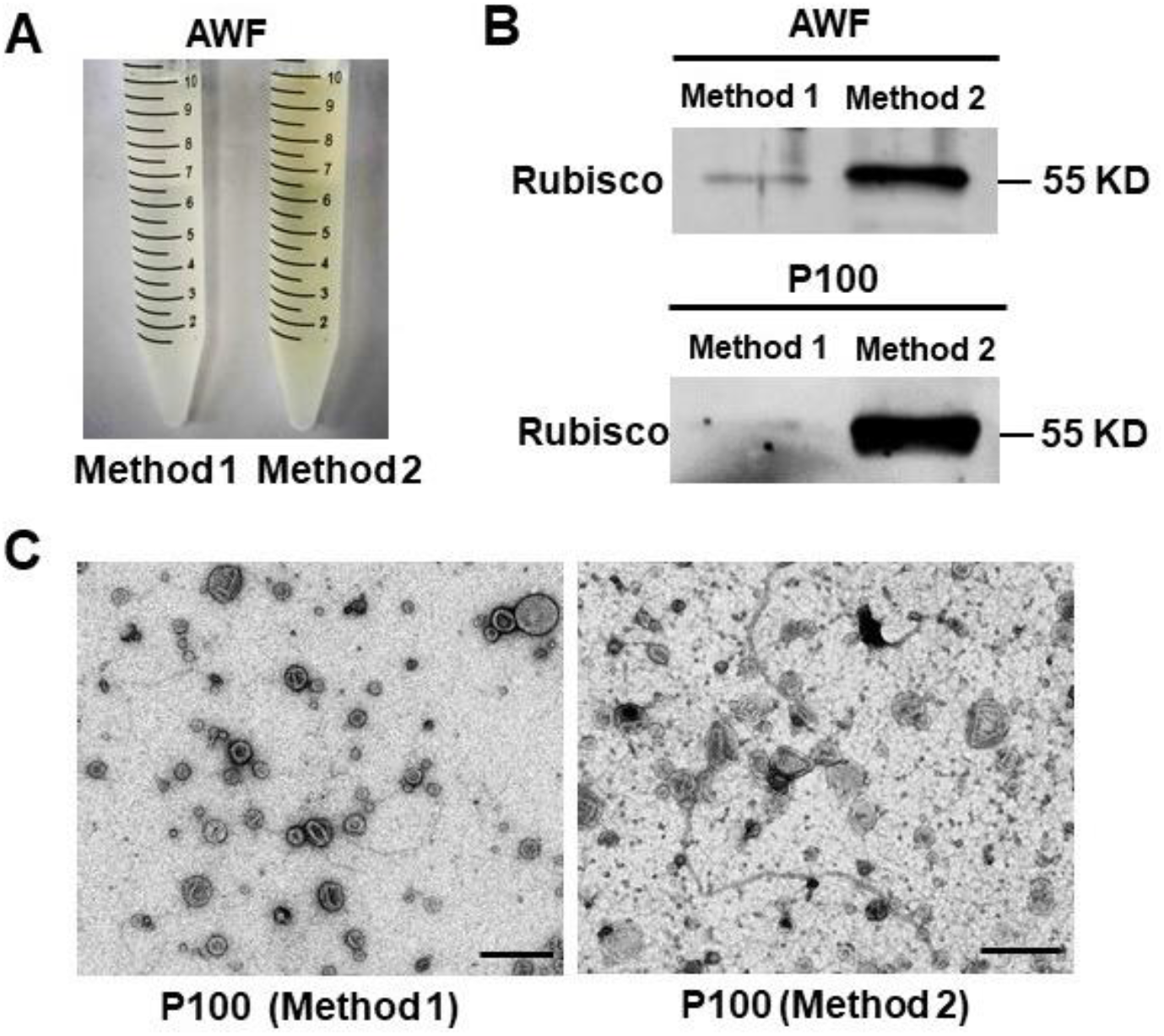
The detached leaves protocol (Method 1) for AWF isolation is better than the whole plant protocol (Method 2) in *Arabidopsis*. (A) Comparison of the color of AWF isolated by Method 1 and Method 2. The same amounts of plants (50 plants) were used for both methods. (B) Detecting Rubisco protein in both AWF and their P100 EV fraction by western blot using Rubisco antibody, the protein size was indicated by KD. To perform the western blot of AWF samples, the same amounts of AWF (10 μl) collected by both methods in (A) were used. To perform the western blot of EV samples, all AWF collected in (A) was centrifuged at 100, 000 **×** g to get P100 fractions. Both P100 pellets were resuspended in 100 μl infiltration buffer, and 10 μl of this suspension was used for the western blot. (C) Representative transmission electron microscopy (TEM) images of P100 fraction isolated from AWF collected by Method 1 and Method 2. Scale bars, 500 nm.

### Technical evaluation of final ultracentrifuge speed for plant isolation

In differential ultracentrifuge steps, the final supernatant is ultracentrifuged to pellet the EVs. In animal systems, genuine exosomes (or small EVs in general) are usually sedimented at speeds of 100,000-200,000 × g (Thery et al. 2006; Kowal et al. 2016; Konoshenko et al. 2018; Jeppesen et al. 2019). For plant EV isolation, two final ultracentrifuge speeds, 100,000 × g (Prado et al. 2014; Prado et al. 2015; Cai et al. 2018; Movahed et al. 2019; Liu et al. 2020; He et al. 2021) and 40,000 × g (Regente et al. 2009; Rutter and Innes 2017; Baldrich et al. 2019), have been used in different studies. Here, we compared the final ultracentrifugation steps, by centrifuging clean AWF at 100,000 × g, or at 40,000 × g, to obtain two EV fractions, named P100 and P40, respectively (Figure 3A). The pellet obtained from further centrifugation of the supernatant of the P40 fraction at 100,000 × g was named the P100 minus P40 (P100-40) fraction (Figure 3A). Subsequently, the morphology of EVs were examined by TEM (Jung and Mun 2018). EVs in the P100 fraction showed similar morphology to animal EVs isolated by centrifugation at 100,000 × g (Figure 3B) (Thery et al. 2006; Jung and Mun 2018). Plant EVs are unlikely to be deformed or broken during 100,000 × g centrifugation (Figure 3B). There were less EVs in the P40 fraction in comparison to the P100 fraction, while a substantial amount of EVs were isolated after centrifugation of the supernatant of fraction P40 at 100,000×g (P100-40) (Figure 3B), therefore, it is wrong to call the supernatant fraction of P40 vesicle-depleted fraction as claimed in Baldrich *et al*., 2019 (Baldrich et al. 2019). Furthermore, similar to the previous conclusion, isolation of EV fractions from transgenic plants co-expressing TET8-GFP and mCherry-PEN1 proteins that could monitor TET8 positive and PEN1 positive EVs, showed a large amount of TET8 positive EVs in the P100-40 fraction (Figure 3C). These results demonstrate that centrifugation at 100,000 × g has higher separation efficiency than centrifugation at 40,000 × g for plant EV isolation, and it results in a loss of large amounts of TET8 positive EVs if using 40,000 × g.

**Figure 3.**
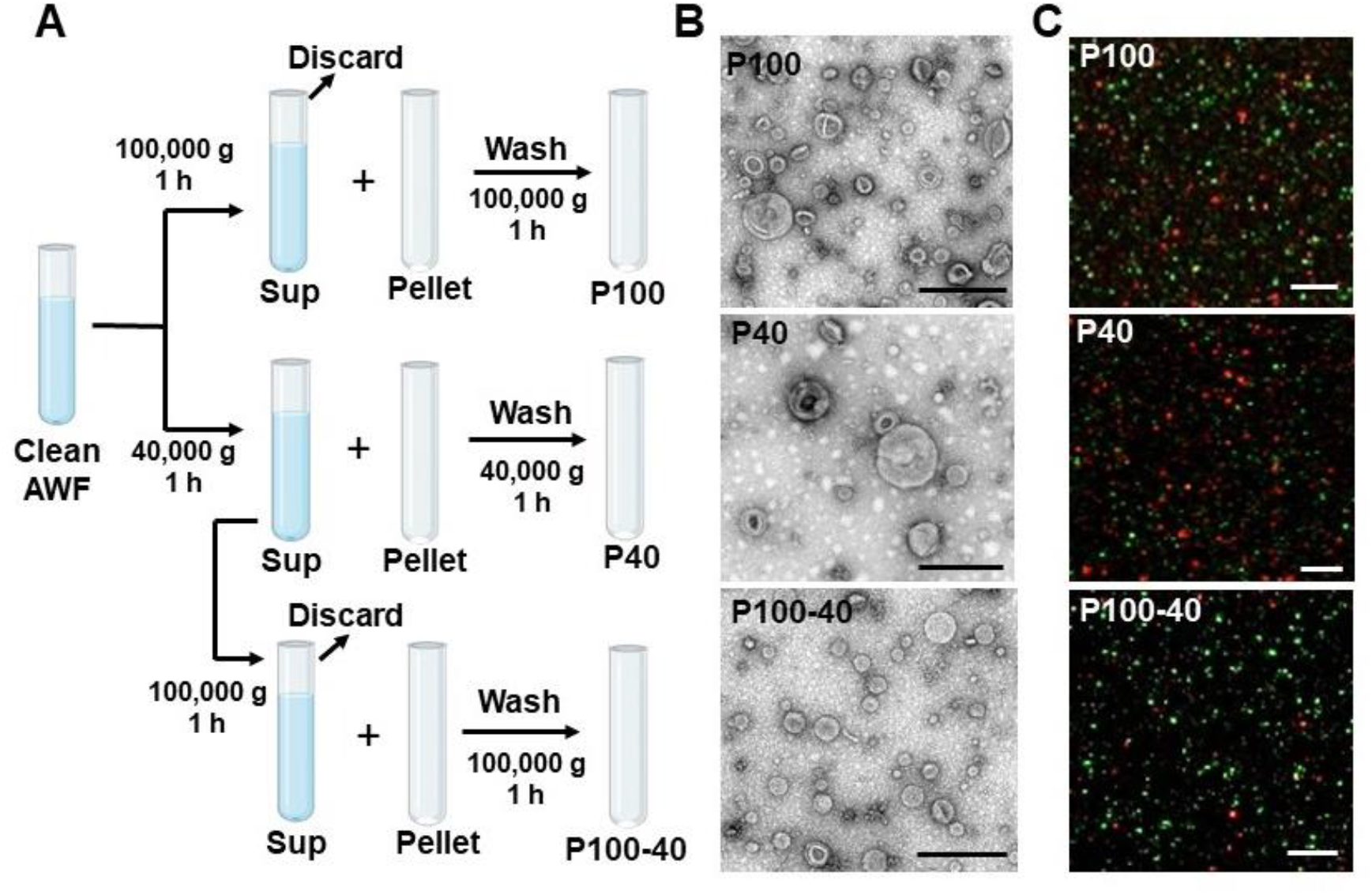
Centrifugation at 100,000 × g enriches plant EVs much more efficiently than at 40,000 × g. (A) Schematic of EV isolation by differential ultracentrifugation of AWF from *Arabidopsis*. EVs isolated from clean AWF (isolated by Method 1) via ultracentrifugation at 40,000 **×** g (P40 fraction) and 100,000 × g (P100 fraction) for 1 hour. For the P100-40 fraction, the supernatant (Sup) of P40 fraction was centrifuged a second time at 100,000 ×g for 1 hour. (B) Representative TEM images of P100 fraction, P40 fraction and P100-40 fraction isolated from *B. cinerea*-infected wild-type *Arabidopsis*. Scale bars, 500 nm. (C) Confocal microscopy of EV fractions (P100, P40 and P100-40) isolated from *B. cinerea*-infected *TET8-GFP/mCherry-PEN1* double-fluorescence transgenic plants. Scale bars, 5 μm.

We also analyzed the plant EV size based on TEM imaging. The TEM micrographs of the P100 fraction showed that a majority of the vesicles (92.3%) had diameters ranging between 30 nm and 150 nm, and an average diameter of 84.5 nm (Figure 4). This result demonstrated that plant EVs in the P100 fraction are similar in size to an EV subtype termed exosomes (30 nm-150 nm in diameter) (Colombo et al. 2014; Kowal et al. 2016; Mathieu et al. 2019). The average size of vesicles observed in the P40 fraction was 97.8 nm, while vesicles in the p100-40 fraction, had an average diameter of 69.5 nm (Figure 4), suggesting that centrifugation at 40,000 × g pellets larger vesicles. These results suggest that centrifugation at 100,000 × g enriches plant EVs, especially small EVs like exosomes, much more efficiently than centrifugation at 40,000 × g.

**Figure 4.**
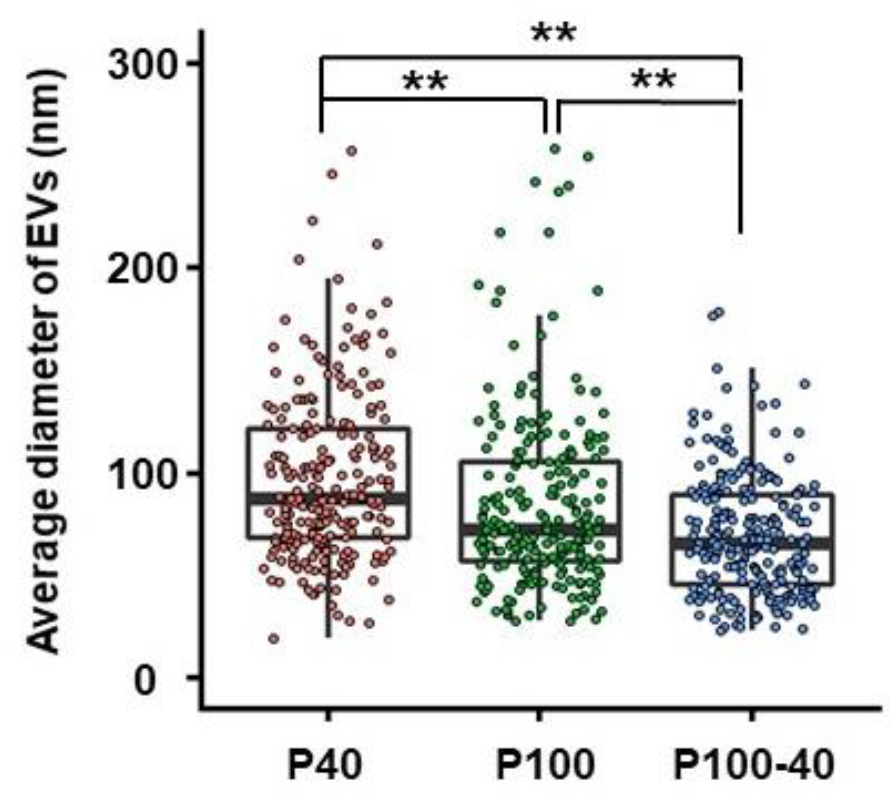
Histograms for the size distribution of EVs in P40 (n = 226 particles analysed, mean = 97.8 nm), P100 (n = 222 particles analyzed, mean = 84.5 nm) and P100-40 (n = 232 particles analyzed, mean = 69.5 nm) fractions from TEM images. The statistical analysis was performed using ANOVA Dunnett’s multiple comparisons test. Each open circle represents a single EV value. ^**^P < 0.01.

### Density gradient fractionation separates plant EVs

Although Method 1 provides reasonably pure plant EVs (Figure 2), some applications may require extra purification steps. The overlapping sedimentation of exosomes, microvesicles and other large vesicles produces a mixture of vesicles in the ultracentrifugation fraction (Konoshenko et al. 2018). Density gradient fractionation separation is a classical method used to separate vesicles according to their floatation speed and equilibrium density (Colombo et al. 2014; Kowal et al. 2016; Jeppesen et al. 2019). This strategy separates EVs using sucrose or iodixanol gradient centrifugation of EV pellets prepared by differential ultracentrifugation. For plant EV isolation, vesicles isolated from the P100 fraction have been floated in a sucrose gradient (He et al. 2021) and vesicles isolated from the P40 fraction have been floated in an iodixanol (OptiPrep) gradient (Rutter and Innes 2017) to facilitate the separation of subtypes of EVs. Because centrifugation at 100,000 × g enriches plant EVs much more efficiently than 40,000 × g, we used an iodixanol density gradient to further separate EVs from the P100 fraction and estimate their density using top-loading methods (Figure 5A). Using the TET8 antibody, we identified that most of the TET8 positive EVs accumulated in the third fraction (F3) at the average density of 1.08 g/ml of iodixanol (Figure 5B). This is similar to the density of exosomes in animal systems (1.08–1.12 g/ml) (Wubbolts et al. 2003; Iliev et al. 2018; Jeppesen et al. 2019). TET-positive EVs (exosomes) are also enriched in F3 fraction of iodixanol gradient. We further analyzed the size of vesicles in the F3 fraction using TEM imaging. The majority of vesicles (93%) in the F3 fraction had diameters ranging between 30 nm and 150 nm, and with an average diameter of 68.9 nm. This is similar to the EV sizes obtained in the P100-40 fraction (Figure 4, Figure 5C and 5D).

**Figure 5.**
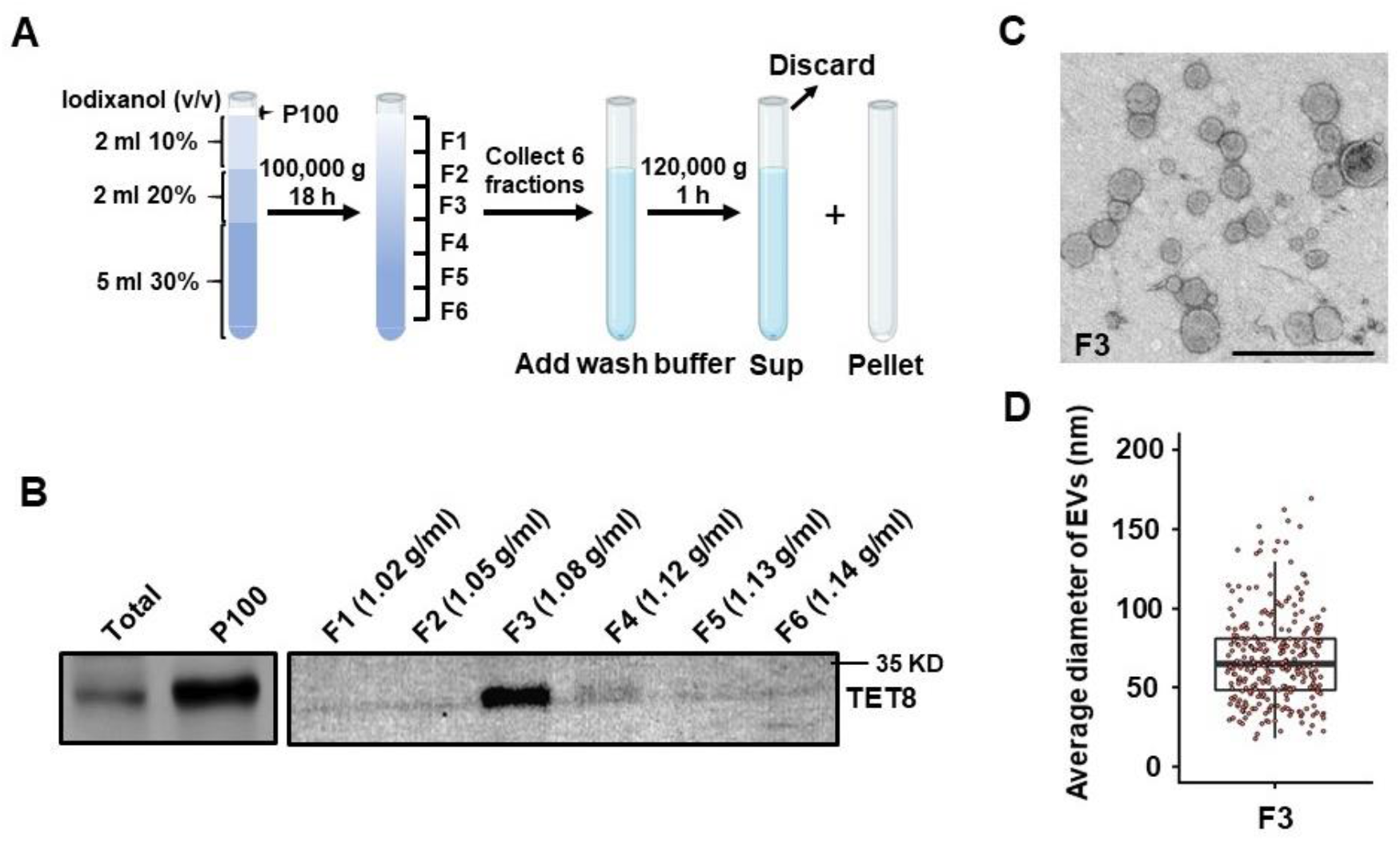
Characterization of P100 EVs using iodixanol gradients. (A) P100 fraction obtained after 100,000× g centrifugations was allowed to float into an overlayed iodixanol gradient by top loading. (B) Six fractions were collected and analyzed by western blot, showing the TET8-positive EV enriched in a single fraction (F3) with TET8 native antibody. (C) Representative TEM images of F3 fraction in (B). Scale bars, 500 nm. (D) Histogram for the size distribution of EVs in F3 fraction in (B) (n = 284 particles analyzed, mean = 68.9 nm). Each open circlerepresents a single EV value.

### Immunoaffinity capture-based technique to isolate plant EVs

Immunoaffinity capture-based technique is considered the most advanced method to purify a specific class of EVs (Thery et al. 2006; Kowal et al. 2016; Jeppesen et al. 2019; He et al. 2021). This technique relies on the use of an antibody to capture EVs based on the expression of a specific marker on the surface of the EVs (Thery et al. 2006). Tetraspanin families such as CD81 or CD63, are ideal immuno-capture molecules since they are enriched on exosome membrane (Andreu and Yanez-Mo 2014). He *et al*. developed an immunocapture purification method using beads coated with an antibody targeting plant exosome marker TET8 (Figure 6A)(He et al. 2021). It is worth noting that antibody-recognized regions of the protein marker must be on the surface outside of EVs. Thus, the antibody that specifically recognizes the large exposed extravesicular loop, EC2 domain of TET8 has been well designed to pull-down TET8 positive EVs from the P100 fraction (He et al. 2021). Using this method, TET8-positive EVs can be successfully isolated from the P100 fraction, and then easily detected by confocal microscopy (Figure 6B). Specificity of the immunoaffinity capture was examined using beads coated with an irrelevant antibody (IgG) (Figure 6B). Thus, this approach can be easily used for isolating/purifying a specific subtype of EVs in plants. By using this method, EV-enriched sRNAs and RNA binding proteins, such as AGO1, RH11, RH37, ANN1 and ANN2 were clearly detected in the TET8-positive EVs (He et al. 2021). Thus, the immunoaffinity isolation should be the best method for the precise analysis of the cargo contents of specific EV subtypes.

**Figure 6.**
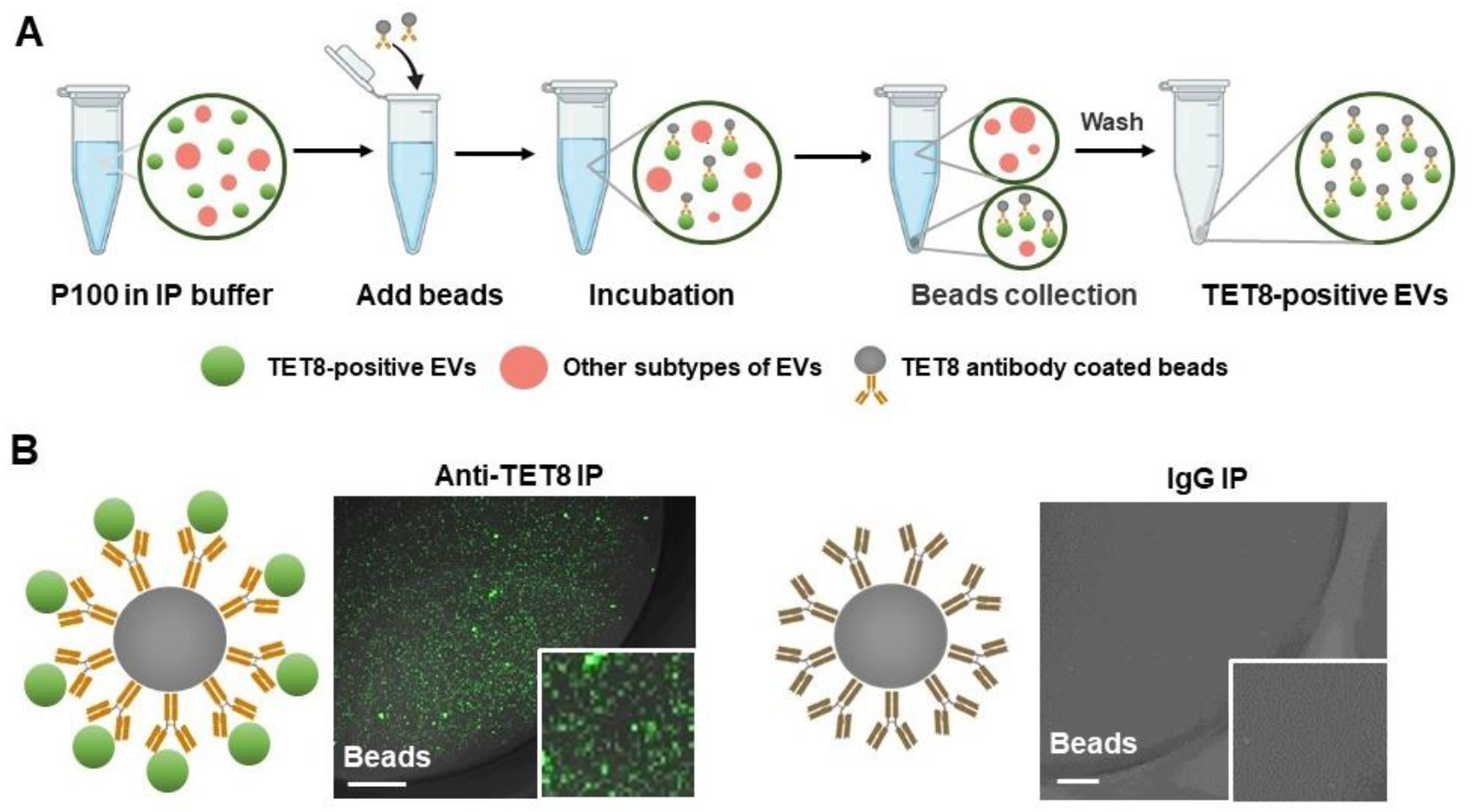
Immunoaffinity isolation is the most advanced method for the purification of specific subclass of EVs in plant. (A) Schematic of P100 fraction was subjected to parallel immuno-isolation with beads coupled to antibodies against TET8, or irrelevant Rabbit IgG as a negative control. (B) Confocal microscopy showed that the TET8-positive EVs pulled-down by TET8-specific antibody-linked beads. Scale bars, 10 μm.

## DISCUSSION

In recent years, plant-derived EVs have gained more and more interest because of their essential role in organism-to-organism communication, and research in this field has exponentially increased (Cai et al. 2019a; Cai et al. 2021). Although in some literature, the nano-sized particles isolated from disrupted leaves or plant tissues by differential ultracentrifugation are also considered plant EVs, they are actually artificial plant-derived vesicles instead of true natural EVs (Zhang et al. 2016a; Kameli et al. 2021). Interestingly, these artificial vesicles have shown promising utility in human health and disease applications (Wang et al. 2013; Mu et al. 2014; Teng et al. 2018; Liu et al. 2020). Due to the existence of diverse EV separation methods developed in plants, in this study, we compared the quality of EVs isolated from two major published methods step-by-step and developed the most optimal protocol for AWF collection and differential ultracentrifugation with minimum contamination and high yield. Extra subsequent steps, such as immunoaffinity technique, were also introduced to purify specific subclass of EVs.

In animal systems, EVs have been isolated from diverse bodily fluids, including blood, urine, saliva, breast milk, semen and cell culture media (Colombo et al. 2014). In plants, the first critical step for EV isolation is the collection of clean apoplastic fluids. Due to the fragility of plant leaves, collection of clean apoplastic fluids has been a challenge. Here, we found that using only the detached *Arabidopsis* rosette leaves with no petioles for AWF collection can minimize the contamination of cytosolic contents and vasculature fluid (Method 1 in Figure 2). Because phloem stream contains large amounts of mobile RNAs and ribonucleoprotein complexes (Zhang et al. 2009; Liu and Chen 2018), this detached leaf protocol removes those irrelevant RNAs and minimizes the impact on subsequent analyses of RNA content in EVs. On the contrary, the other AWF collecting method using the whole aerial part of a plant (Method 2 in Figure 2), can lead to cell breakage and contamination of intracellular contents and phloem sap. Therefore, isolating AWF from detached leaves with no petioles is more suitable as the first step of plant EV isolation to minimize contamination.

Ultracentrifugation remains the most commonly used technique for EV isolation from different biofluids and cell culture supernatants. Similar to animal EV separations, which overwhelmingly relied on 100,000 × g fractions, plant EVs and their RNA contents were highly enriched in the fraction collected by ultracentrifugation at 100,000 × g from AWF (Cai et al. 2018; He et al. 2021). In this study, we have characterized EVs isolated from both the intermediate speed (40,000 × g) or high speed (100,000 × g) fractions. Consistent with our previous study (He et al. 2021), we found that centrifugation at 100,000 × g has higher separation efficiency resulting in higher EV yield, whereas centrifugation at 40,000 × g largely reduces the yield, and tends to collect large EVs. There is still large amount of EVs, especially TET-positive exosomes, in the P100-40 fraction. When using AWF collected by Method 2, centrifugation at lower speeds, such as 40,000 × g, may be suitable for isolating PEN1-associated EVs (Rutter and Innes 2017; He et al. 2021), we also observed a large amount of non-vesicle material impurities in the P100 fraction, further indicating the bad quality of AWF collected from whole aerial plant material by Method 2.

The density gradient centrifugation technique enables the isolation of EVs with higher purity and the separation of distinct EV subtypes (Konoshenko et al. 2018; Jeppesen et al. 2019). Previously, we found that in the P100 fraction floated in a sucrose density gradient, TET8-positive EVs and EV-enriched sRNAs were enriched in the EV fractions at densities of 1.12-1.19 g/ml (He et al. 2021). In this study, P100 fractions were floated in an iodixanol density gradient, and TET8-positive vesicles were enriched in the gradient fraction of 1.08 g/ml, on average. The different density of TET8 positive EVs in sucrose versus iodixanol could be a result of differences in the osmotic pressure of these two gradients. This result was similar to a previous study that showed the difference in separation of P100 pellets derived from human dendritic cells in sucrose versus iodixanol gradients (Kowal et al. 2016). Note that PEN1-positive EVs collected at 40, 000 × g were enriched in the iodixanol gradient fraction at densities ranging from1.029 g/ml to 1.056 g/ml (Rutter and Innes 2017), supporting that TET8-positive EVs and PEN1-positive EVs are two different sub-populations of EVs with distinct densities. Further study is required to determine the density of other EV subtypes, such as Exo70E2 positive EVs, by its marker lines or the specific antibodies.

Density gradient centrifugation still has some disadvantages, as it is complex, laborious, and time-consuming (up to 2 days). Additionally, it is difficult to separate different subtypes of EVs with similar densities. Immunoaffinity isolation is the most precise method for purifying a specific subtype of EVs (Thery et al. 2006; Kowal et al. 2016; Jeppesen et al. 2019; He et al. 2021). Co-isolation of non-vesicular contaminants from the cytoplasm and other unwanted vesicle can be prevented by the highly specific affinity interactions that occur between an antigen and an antibody. The antigens ideally are EV biomarkers that are highly concentrated on the EV membranes, for example, the Major histocompatibility complex (MHC) antigens and tetraspanins proteins on exosomes (Kowal et al. 2016; Jeppesen et al. 2019). In plants, we showed that TET8-positive EVs can be successfully isolated from P100 fraction by an antibody that specifically recognizes the EC2 domain of TET8. It would be ideal to purify other subclass of plant EVs using immunoaffinity isolation method. In summary, these findings should serve as a guide to choose and further optimize EV isolation methods in the plant field for their desired downstream applications.

## MATERIALS AND METHODS

### Plant materials

*Arabidopsis thaliana* ecotype Columbia-0 (*Col 0*) was used in this study. *Arabidopsis* marker lines *TET8*_*pro*_*::TET8-GFP* (Cai et al. 2018; He et al. 2021) and *TET8-GFP/mCherry-PEN*1 double-fluorescence lines (Cai et al. 2018; He et al. 2021), were used as described previously. *Arabidopsis* seeds were grown in soil side-by-side at 22 °C for 4 weeks under short-day periods (12 h of light followed by 12 h of darkness).

### Apoplastic washing fluids collection from *Arabidopsis* leaves

Apoplastic washing fluid (AFW) collection from *Arabidopsis* leaves was modified from previous studies (O’Leary et al. 2014; Madsen et al. 2016). A typical experiment for EV isolation requires ∼50 plants for each genotype/treatment. Distinct proximal (petiole) part of leaves was removed by using scissors, and the distal (blade) zones of leaves were collected. After recording the biomass, leaves were washed 3 times with water. The leaves were carefully placed in a 200 ml syringe and gently vacuumed with an infiltration buffer (20 mM MES hydrate, 2 mM CaCl_2_, 0.1 M NaCl, pH 6.0) for 20 seconds. Excess infiltration buffer on the leaf surface were removed by a clean paper towel and then fixed leaves on a small plastic stick. Small plastic stick with leaves were put into a 50 ml conical tube, keeping all leaf opex up, and then centrifuged for 10 min at 4 °C at 900 × g to collect the AWF.

### Isolation of plant EVs by differential ultracentrifugation

Plant EVs were isolated from *Arabidopsis* AWF. The AWF was centrifuged for 30 min at 4 °C at 2,000 × g to remove large cell debris and then filtered by a 0.45 μm filter. Next, the supernatants were transferred into new ultracentrifuge tubes and centrifuges for 30 min at 4 °C at 10,000 × g. After the pellet was discarded, the supernatants (the clean apoplastic washing fluid) were centrifuged for 1 hour at 4 °C at 100,000 × g or 40,000 × g to obtain the P100 EV fraction or P40 EV fraction. To obtain the P100-P40 EV fraction, the supernatants of P40 were centrifuged for 1 hour at 4 °C at 100,000 × g. All pellets were washed in 10 ml of infiltration buffer and finally re-centrifuged at the same speed before being resuspended in infiltration buffer for further study.

### Iodixanol gradient separation of plant EVs

Discontinuous iodixanol gradients (OptiPrep, STEMCELL) were prepared as described in previous protocol with slight modification (Kowal et al. 2016). Working solutions of 10% (v/v), 20% (v/v) and 30% (v/v) idoixanol were made by diluting an aqueous 60% OptiPrep stock solution in infiltration buffer (20 mM MES hydrate, 2 mM CaCl_2_, 0.1 M NaCl, pH 6.0). The gradient was formed by successively layering 4.8 mL of 30% solution, 2.1 mL of 20% solution, and 2 mL of 10% solution in 13PA tube (himac) from bottom to top. About 0.4 mL of EVs were resuspended in infiltration buffer was layered on top of the gradient. The tube was centrifuged for 100,000 × g for 17 h at 4°C (P40ST, CP80NX, himac). After stopping the centrifuge without breaks, 6 fractions of 1.4 ml were collected from top of the tube. These fractions were each brought up to 12 ml with an infiltration buffer and centrifuged at 100,000g for 60 min at 4°C to obtain pellets in each fraction.

### Transmission electron microscope analysis of plant EVs

Sample preparation of EVs for TEM observation referred to Maroto *et al*.(Maroto et al. 2017). 10 μl of EVs suspensed in infiltration buffer was deposited on 3.0 mm copper Formvar-carbon-coated electron microscopy grids (TED PELLA), and then the sample was wicked off using filter paper, and the grids were negatively stained with 10 μl of 1% uranyl acetate. The grids were allowed to air dry and imaged at 100 KV using Transmission Electron Microscope (JEM-1400plus, JEOL). EV size was assessed with Image J software.

### Immunoaffinity capture of plant EVs

Immunoaffinity capture of plant EVs was performed as described previously (He et al. 2021). Briefly, antibodies for immunoaffinity capture, Rabbit polyclonal anti-AtTET8 (Homemade) and normal rabbit immunoglobulin G (Thermo Fisher), were coated with protein A beads in IP buffer (20 mM MES hydrate, 2 mM CaCl_2_, 0.1 M NaCl, pH 7.5). Beads were then washed 3 times with an IP buffer (containing 0.3% BSA), and resuspended in the same buffer, to which P100 fraction was added, followed by overnight incubation at 4 °C with rotation. Bead-bound EVs were collected and washed by IP buffer for further analysis.

### Confocal microscopy analyses of plant EVs

For visualization of EV-associated GFP-fluorescence and mcherry-fluorescence, EV pellets or EV coated beads were suspended in infiltration buffer were examined using a 40x water immersion or dip-in lens mounted on a Confocal Laser Scanning Microscope equipped with an argon/krypton laser (Leica TCS SP5).

## ACKNOWLEDGEMENTS

Work in the Q.C. laboratory was supported by grants from the National Natural Science Foundation of China (32070288), and supported by Hubei Hongshan Laboratory. Work in the H.J. laboratory was supported by grants from the National Institute of Health (R35 GM136379), the National Science Foundation (IOS2017314), the United States Department of Agriculture National Institute of Food and Agriculture (2021-67013-34258 and. 2019-70016-29067), the Australian Research Council Industrial Transformation Research Hub (IH190100022), as well as the CIFAR Fungal Kingdom fellowship to H.J.

## AUTHOR CONTRIBUTIONS

H.J. and Q.C. designed the experiments. Y.H. and S.W. performed the experiments. Y.H. and Q.C. analyzed the data. Q.C. and H.J. wrote the manuscript. All authors read and approved of its content.

---

**Figure S1.**
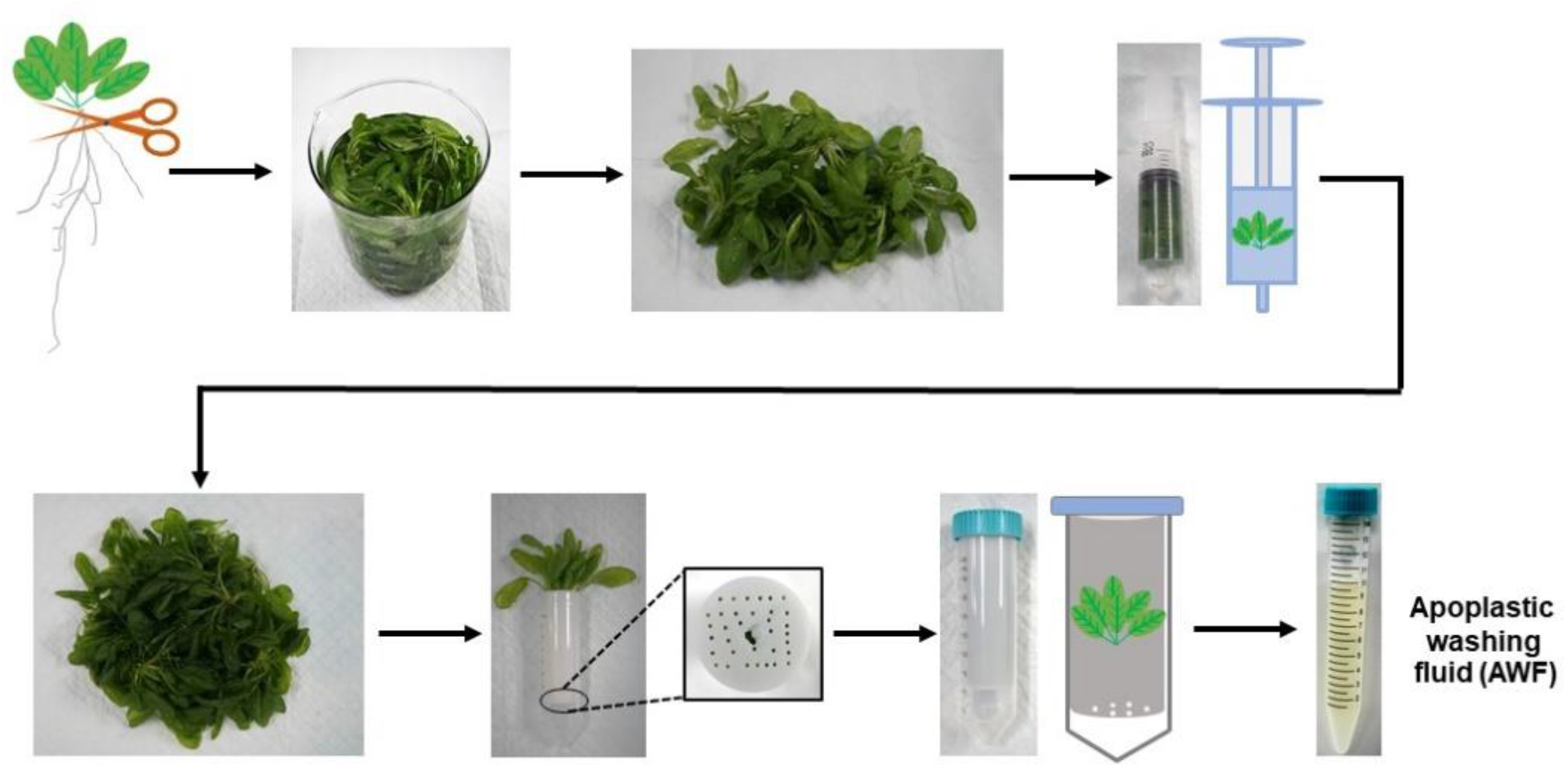
Work-flow of isolation of AWF from *Arabidopsis* (Whole rosettes protocol, Method 2 in Figure 2). Whole rosettes were harvested at root by using scissors. The rosettes were placed in a syringe and gently vacuumed with infiltration buffer, and then placed root down into a 30 ml tube, which was then put into 50 ml conical tube, and then centrifuged at 900 × g to collect the AWF.

